# Phase of neuronal activity encodes 2-dimensional space in the human entorhinal cortex

**DOI:** 10.1101/352815

**Authors:** Zoltan Nadasdy, Ágoston Török, T. Peter Nguyen, Jason Y. Shen, Deborah E. Briggs, Pradeep N. Modur, Robert J. Buchanan

## Abstract

The entorhinal cortex plays a vital role in our spatial awareness. Much focus has been placed on the spatial activity of its individual neurons, which fire in a grid-like pattern across an environment^1^. On a population level, however, neurons in the entorhinal cortex also display coherent rhythmic activity known as local field potential. These local field oscillations have been shown to correlate with behavioural states but it remains unclear how these oscillations relate to spatial behaviour and the spatial firing pattern of individual neurons. To investigate this, we recorded entorhinal cortical neurons in the human brain during spatial memory tasks performed in virtual environments. We observed a spatial modulation of the phase of action potentials relative to the local field potentials. In addition, the spike phase modulation displayed correlation with the movement of the avatar, displayed discrete phase tuning at the cellular level, rotated phase between electrodes, and expressed spatially coherent phase maps that scaled with the virtual environment. Using surrogate data, we demonstrated that spike phase coherence is dependent on the spatial phase dynamics of gamma oscillations. We argue that the spatial coordination of spike generation with gamma rhythm underlies the emergence of grid cell activity in the entorhinal cortex. These results shed a new light on the intricate interlacing between the spiking activity of neurons and local field oscillations in the brain.

---

The medial entorhinal cortex (EC) is found in the medial temporal lobe of the mammalian brain and plays a critical role in spatial awareness in rodents^1^ as well as in humans^2–4^. The activity of grid cells and border cells in the EC, together with place cells and head direction cells in the hippocampus and subiculum, respectively, constitute key components of the allocentric navigation system enabling individuals to localize themselves relative to the environment. The spatially periodic activity, a hallmark of grid cells^1^ and displayed by about 50% in the human brain^4^, is hypothesized to support spatial navigation as an internal coordinate system.

In addition to their spatial tuning, EC neurons generate a broad frequency range of local field potentials (LFPs) with two prominent harmonic components: theta and gamma^5^. Theta and gamma are predominant in the rodent and human hippocampus and EC during active exploratory behaviour and REM-sleep^2,6^ and hippocampal theta tend to phase-couple with slow gamma (30-50 Hz) from EC^7,8^. Theta-spike and theta-gamma phase coherence are instrumental for memory encoding^9,10^. Although grid cells represent the allocentric coordinate system for spatial navigation in a number of mammalian species^1,11,12^, the role of oscillations in the generation of this spatially periodic activity is unknown.

A grid of 16 extracellular microelectrodes (Fig. 3a) was implanted in the EC of two human subjects for seizure monitoring. We analysed the single-unit activity (Fig. 1a) of 525 neurons with sufficiently large inter-cluster Mahalanobis distances and simultaneous LFPs recorded from a separate electrode (Fig. 1b,e) during spatial navigation task in 4 virtual environments (total 2100 data epochs). The spike phase analysis was focused on the two most prominent frequency bands of LFP significantly deviating from the 1/f function, 2-12 Hz (theta) and 25-35 Hz (gamma) (Fig. 1b)^6,13^. We reasoned that if grid cells fire in spatially periodic locations while maintaining phase-locking with LFP then the 2D spatial projection of spike phases is expected to unravel an allocentric spatially periodic pattern.

**Figure 1:**
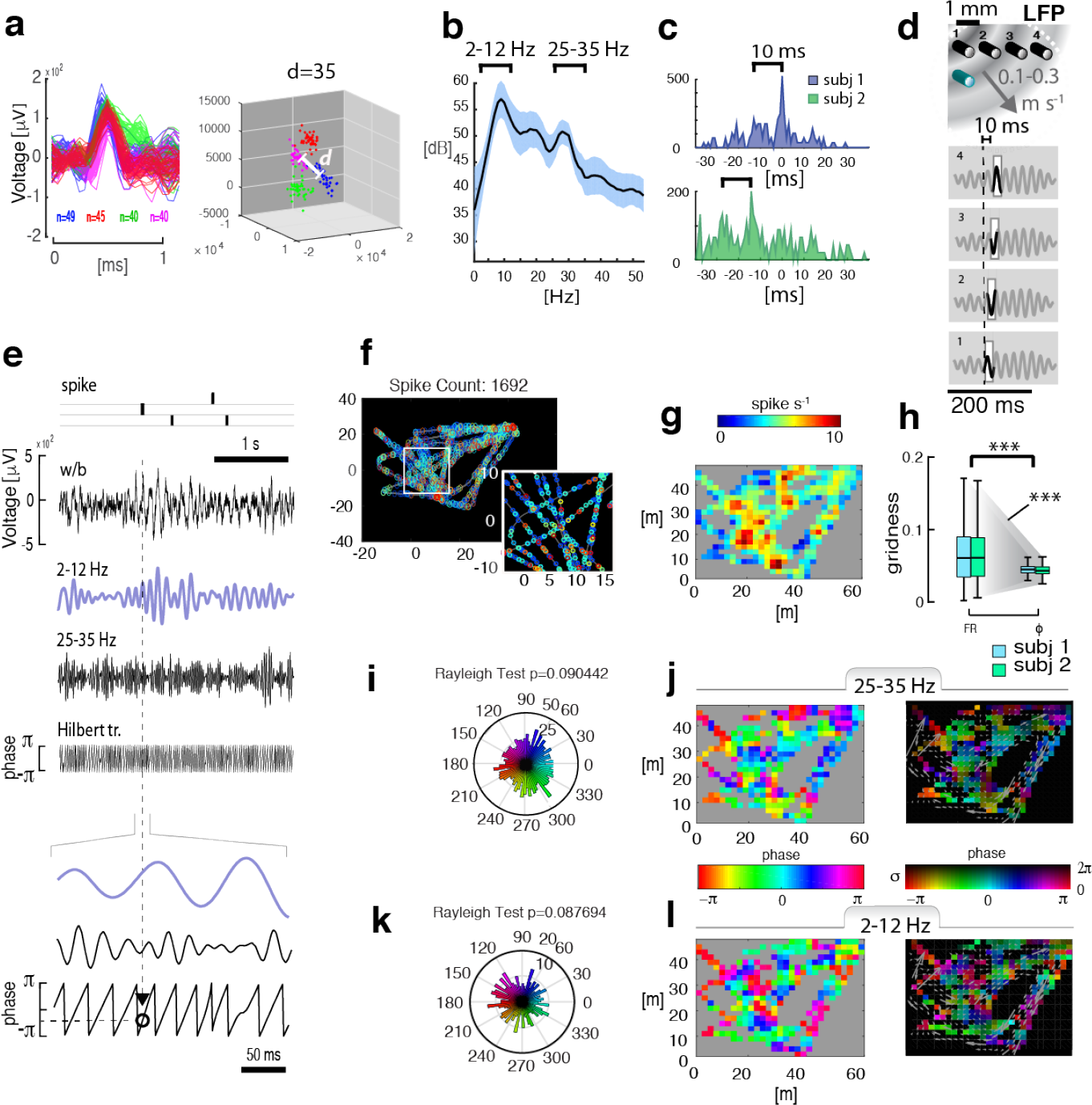
Converting single-unit activity to gamma and theta phase maps. (a) Spike waveforms of four single-units recorded from one of the 16 electrodes and corresponding waveform clusters (20 min data). The two clusters with the shortest Mahalanobis distance between them is indicated. (b) Average spectral density of LFP sampled from a separate electrode during the same 20 min spatial memory task. Braces represent the most prominent frequency components at theta (2-12 Hz) and gamma band (25-35 Hz) band before filtering. Shaded area is SEM. (c) Inter-electrode cross-correlograms of gamma LFP peaks combined across electrode pairs from two subjects (5 min recording). (d) The schematic of the propagating gamma field relative to electrodes (green pin represents the electrode devoted to LFP). The constant delay between the LFP peaks (panel 1-4) sampled by different microelectrodes is consistent with the periodic pattern of cross-correlograms in (c). (e) Computing phase of spikes relative to LFP illustrated on 3 s data. From the top: Spike times of 3 neurons; broad-band LFP (0.1-300 Hz); band-pass filtered LFP at theta (2-12 Hz) and gamma (25-35 Hz); the Hilbert-transform of gamma; theta, gamma and the Hilbert-transform of gamma magnified. The point of intersection between spike time and the Hilbert-transform of gamma (or theta) defines the spike phase. (f) Paths traveled by the avatar during a 5 min navigation task in the Louvre environment. The colour of symbols represent single-units of different neurons. Inset shows a magnified view. (g) The map of mean firing rate per *m*^2^ overlaid the area of the environment. (h) Comparisons of gridness scores between firing rates and gamma-phases combined from two subjects (blue and green represent subject 1 and 2, respectively). (i) Angular histogram of spike phases (*ϕ*) relative to gamma cycles. (j) The spatial distribution of average phases of spikes (Φ) relative to gamma after spatial binning per *m*^2^. (k) Angular histogram of phase ditribution of *ϕ* relative to theta (axes as in i). (l) The spatial distribution of Φ relative to theta per *m*^2^. Colourscale in (j) and (l) represent average phase from − *π* to *π* and hue represents variance of phase (darker is larger). Each pixel covers one *m*^2^ of space of the virtual environment (Louvre in this example). Small gray arrows overlaid are resultant vectors of heading direction. velocity.

We computed the instantaneous phase of each spike event (*ϕ*) using the Hilbert transform of theta and gamma. While the distributions of spike phases for most cells differed from uniform, 3-times as many (19%) neurons expressed phase non-uniformity relative to gamma as to theta (6.21%) (Figs. 1i,k, 3a-f, 4a-e; *Rayleigh*_(*gamma*)_*p* < 0.05: 94/478 cells or 376/1914 epochs;binomial test *p*=2.3066e-43 and *Rayleigk*_(*theta*)_*p* < 0.05: 23/370 cells or 92/1480 epochs; binomial test *p*=0.0274) and 10-times as many *ϕ* with *Rayleigh*_(*gamma*)_*p* < 0.01 (Extended Data Fig. 5e). Moreover, the average *ϕ*, examined individually or combined across electrode 1-4, distributed uniformly with respect to theta but displayed significant polarity relative to gamma (Fig. 3a-f forth column and Extended Data Fig. 8; Rayleigh p indicated above circular histograms).

To elucidate the spatial projection of *ϕ*, we divided the area of each environment to size-proportionate units and computed the circular mean and circular variance of *ϕ* within visited unit areas (Methods). By integrating *ϕ* over those areas, we computed a variance-weighted spatial distribution of phases (phase map) denoted by Φ. In addition, we computed firing rate (FR) maps, 2D autocorrelograms of phase maps (Extended Data Fig. 3a-c), the maps of mean heading direction, the resultant vector length (RVL) of the heading direction, and the correlations of Φ with heading direction as well as with RVL from daily navigation sessions in four different environments (Fig. 2a-e). The Φ revealed spatially consistent and periodically organized map of spike phases with smooth transitions between iso-phase nodes despite the large variance of trajectories in each environment (Extended Data Fig. 9). The patterns of iso-phase nodes resembled those of grid-cell firing, but the cell’s phase-gridness was independent of FR gridness (Figs. 1g,j and 2a,c). Moreover, the variance of grid scores computed from the phase of all cells (Methods) was significantly smaller than grid scores’ computed from FR maps (coefficient of variation *CV*_*FR−gridmod*_ = .5997 and *CV*_*phase−gridmod*_ = .2623 and two-way ANOVA *F*_*subject*_ = 0.73 *p*_(1,2094)_ = 0.3917 *F*_*FR,ϕ*_ = 143.02 *p*_(1,2094)_ = 0.00097; Fig. 1h).

To unravel the spatial pattern of Φ, despite the average low firing rate of cells (<1Hz; Fig. 4b-d), we combined the activity of 20 single-units during a 5-minute epoch and computed the average phase map and the phase maps of isolated single-units separately. The angular distribution of gamma *ϕ*, when combined across all electrodes in a given environment, was always uniform (Extended Data Fig. 3a), while the same set of neurons displayed a non-uniform spatial distribution of Φ when projected in the space of virtual navigation (Figs. 1j,2c, and Extended Data Fig. 3a).

**Figure 2:**
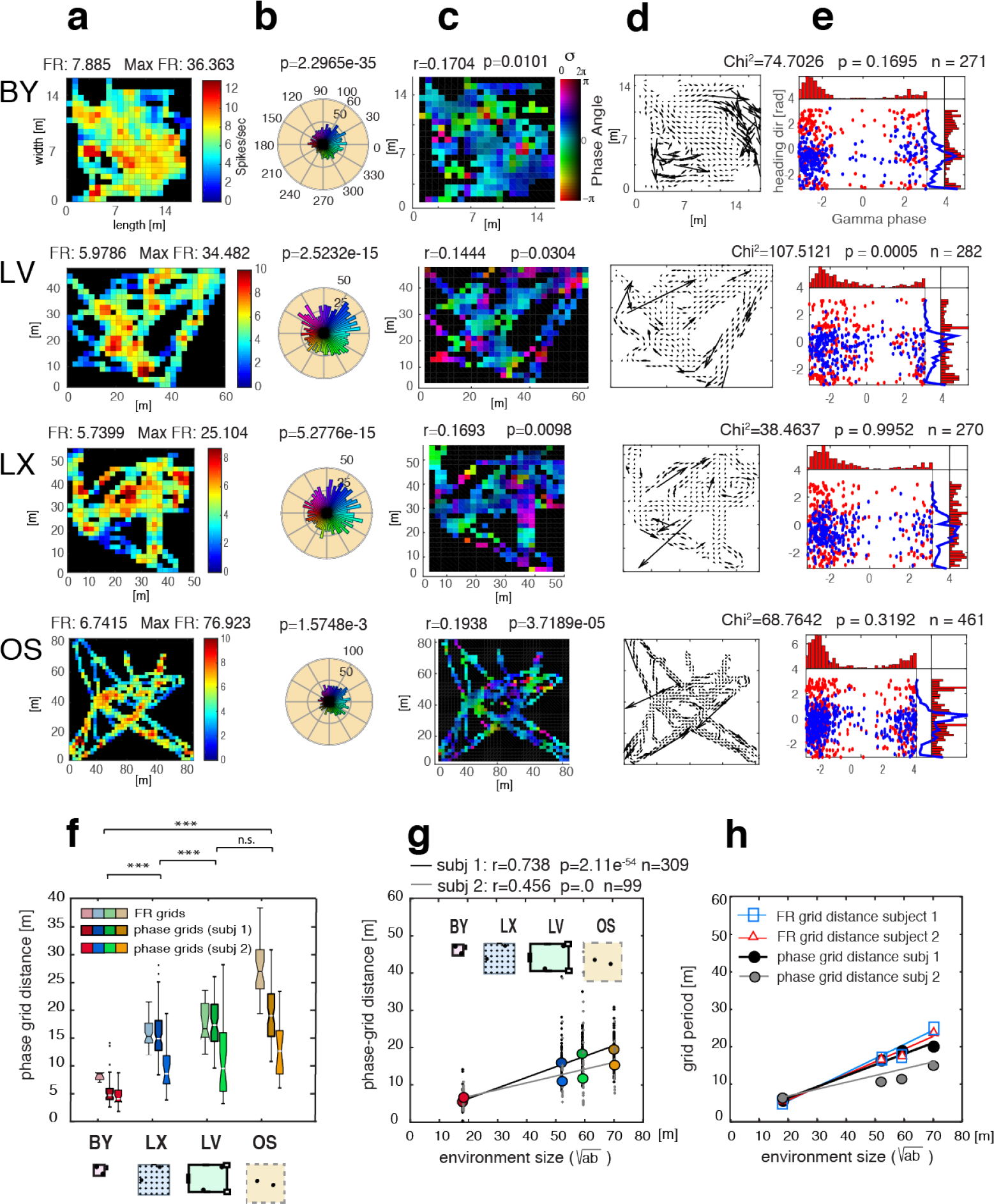
Environment dependent scaling of phase maps. (a) Firing rate maps of the spatial activity of a combined set of four single-units monitored across four different environments (BY,LV,LX,OS). The axis scaling in meter reflects the dimensions of virtual environments. (b) Polar histograms of gamma phase tuning in the four environments. P values represent the significance of Rayleigh tests for circular non-uniformity. (c) Gamma phase maps weighted by variance (described in Fig. 1). The r and p values represent correlations of gamma phase with resultant vector length (RVL) of heading and their significance, respectively. (d) Local RVL of heading directions. Larger arrows represent less variance in local heading direction. (e) Heading direction and gamma phase covariance. Red symbols depict RVL direction per unit area (−*π* < *dir* < *π*) as a function of *ϕ*, blue symbols depict the z-transform of RVL as a function of *ϕ*. Histograms capture the distributions of symbols projected to the phase angle (x red), and to heading direction (y red) and RVL (y blue). χ^2^ and p values indicate the level of independence between phase and RVL. (f) Comparison of FR-grid and phase-grid scaling across environments. The box-triplets from left to right represent the distribution of mean distances between firing rate grids, and mean distances between iso-phase nodes for the two subjects (boxes represent median, 25th and 75th percentile of data). The scheme under the chart depicts scale-proportional layouts of environments. (g) Phase grid scaling as a function of environment size in meters (subjects 1 and 2). Pearson’s correlation coefficients between iso-phase-node distance and environment sizes are indicated. (h) Comparison of slopes of environment dependent scaling of phase-grids relative to FR grids.

**Figure 3:**
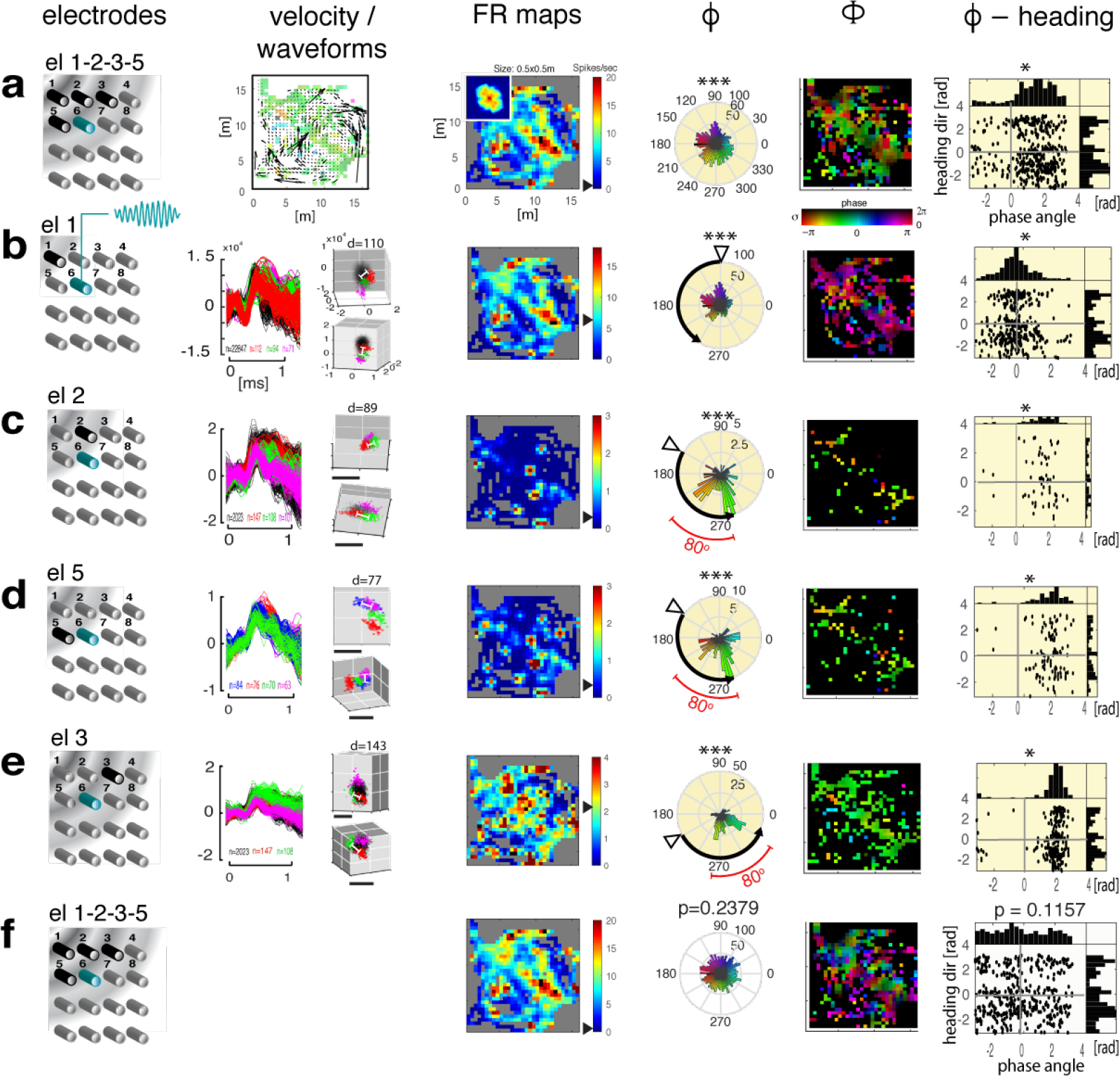
Dependence of Gamma and Theta phases on electrode positions. (a-e) Spatial maps of firing rates and gamma phases of single-units isolated from different electrodes. (f) Spatial maps of single-unit activity relative to theta LFP. First column: 3D schemes of the spatial configuration of microwire electrodes. Gray-shading represent estimated gamma phase difference between electrodes. Second column: Average spatial velocity map of 5 min navigation computed over 0.7 × 0.7 m areas. Second column B-E: single-unit activity (4 cells per electrode) isolated from different electrodes. Waveforms of spikes and corresponding clusters. The smallest of the Mahalanobis distances between cluster centroids is indicated as d. Different single-unit clusters are colour-coded. Third column: Spatial firing-rate maps of single-unit activity isolated then combined from microwires 1-4. Small inset in (a) is the spatial autocorrelogram. Black arrowheads on scalebars represent average firing rates. Forth column: Polar histograms of *ϕ* relative to gamma LFP. Fifth column: Spatial distribution of phases (Φ) relative to gamma. Colours represent phase and darkness represent variance. Spatial dimension of pixels is 0.7 × 0.7 m. Sixth column: The heading direction and gamma phase correlation. Y axis average heading direction associated with the average spike phase within the same unit area (X). Histograms above scatter plots represent phase distribution while histograms at the right sides represent distributions of heading directions. The yellow shading represent electrode at which the correlation between RVL and Φ was significant (p<0.05) (f) Same as (a) except all phases were computed relative to theta LFP. Star symbols represent levels of statistical significance (* p<0.05, *** p<=0.001). Asterisks: above *ϕ* indicate the sigificance of Rayleigh test of directionality; under *ϕ*-heading represent χ^2^ test of phase RVL covariance. The radial histograms were binned by 5 deg, phase maps and average velocity maps were binned by 0.7 × 0.7 m bins.

**Figure 4:**
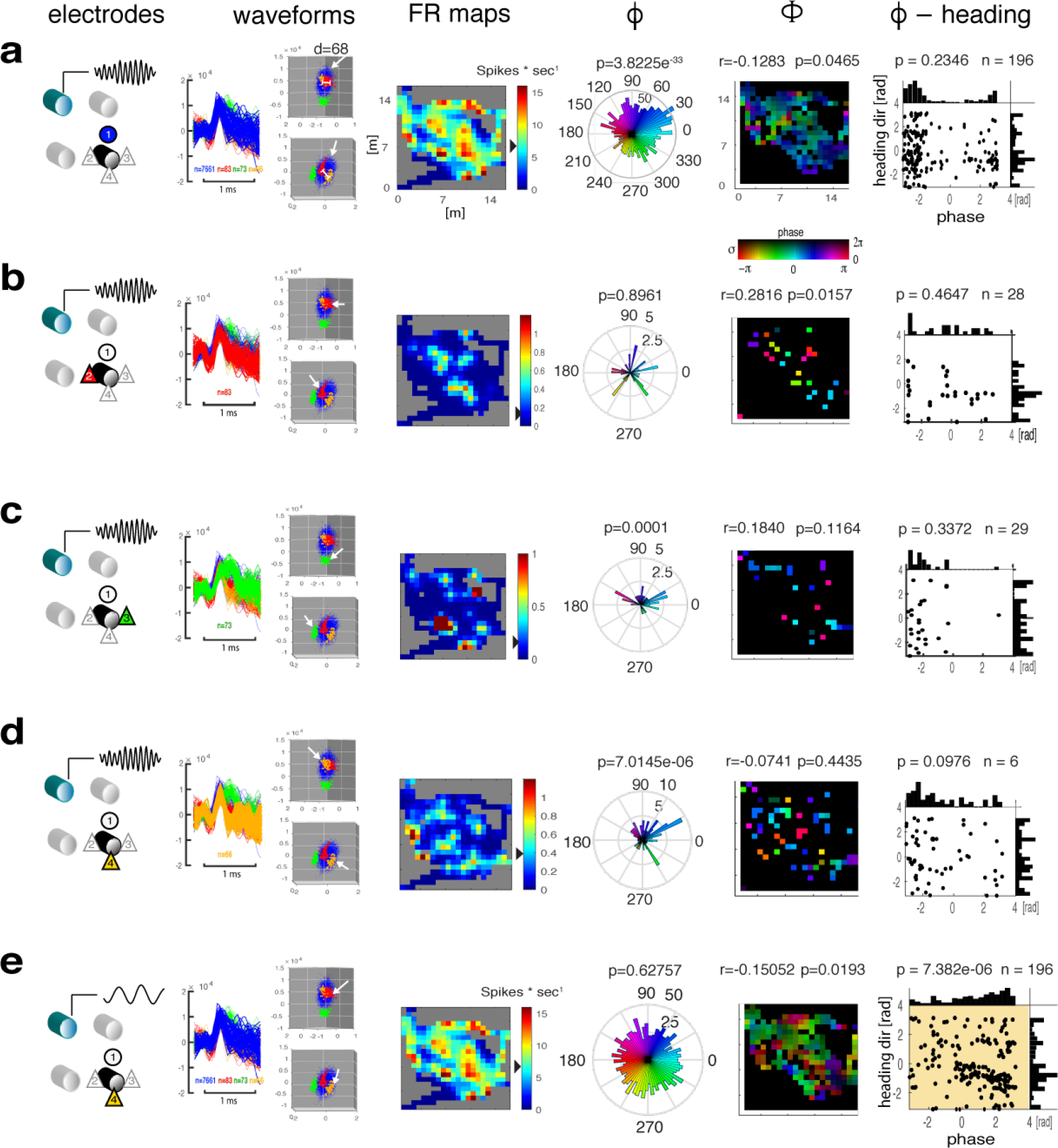
Differences across single-units isolated from the same electrode with respect to gamma and theta phases. (a-d) Spatial maps of firing rates and LFP phases of different single-units isolated from the same electrode. LFP was recorded from a different electrode and filtered at gamma (a-d) and theta band (e) before spike phases were computed. First column: 3D schemes of single neurons recorded from one of the four microwire electrodes and gamma/theta LFP was extracted from a separate distant electrode. Insets: 500 ms LFP samples of gamma (a-d) and theta (d). Second to fifth columns: Combined single-unit activity isolated from one electrode. Spike waveforms and corresponding clusters (marked by white arrows) in feature space. The smallest Mahalanobis distance (d) is indicated above the cluster plot. Different single-unit clusters are colour-coded. Third column: Spatial firing-rate maps of single units (a-D). Black arrowheads on scale-bars represent average firing rates. Forth to sixth columns: Same as in Fig. 3 except *ϕ*, Φ and phase-heading direction correlation plots represent activity of indicvidual cells isolated from the same electrode. Colour and darkness in Φ represent phase and variance, respectively, as in Fig. 1-3). Spatial dimension of pixels is 0.7 × 0.7 m. (e) Same cell as (a) except the average phase plots were computed relative to theta LFP. P values of polar histograms under *ϕ* represent Rayleigh test of directionality and p values under phase-velocity correlations in the sixth column represent χ^2^ test of inhomogeneity. Radial histograms and spatial maps were binned as in Fig. 3.

To test the environmental dependence of Φ, we computed the average gamma iso-phase grid distances for each environment across days and found a monotonic increase with the size of environments (Fig. 2f-h; BY=5.462 m LX=15.812 m LV=18.242 m OS=19.119 m). Even though the FR grids did not co-register with iso-phase nodes, the average distance of iso-phase grids and its scaling with the size of the environment were consistent with the grid scaling computed from FR grids (Fig. 2f,h)^4^. The iso-phase grid distances scaled linearly with the sizes of environments 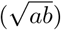 (Fig. 2g; subject 1: R-square=0.5448,RMSE=4.711, f(x)=0.2835x+0.669; subject 2: R-square=0.2026, RMSE=7.395, f(x)=0.1834x+3.087). The slope of iso-phase grid scaling (0.2835 and 0.1834) was slightly smaller but comparable to the slope of firing rate grids (subj-1 = 0.4108 and subj-2 = 0.3509 (Fig. 2h)^4^.

The stability of Φ maps over time and their robustness to spike count variation was validated with two independent cross-validation methods applied to 5 minute epochs (Methods and Extended Data Figs. 4a,b-d and 4a,e-g). The correlations confirmed both stability and robustness to spike sampling of phase maps (Pearson’s *r*=0.438 *p*_(1,359)_=<0.0001 and *r*=0.239 *p*_(1,359)_ <0.0001 for stability and robustness, respectively).

To test the statistical significance of spatial organization of Φ against by-chance pattern-formation and to elucidate the feature that controls the topography of phase maps, we applied two types of surrogate tests: inter-spike-interval permutation, and phase-permutation of LFP (Extended Data Fig. 5a-c). These surrogates retained the average firing rate of neurons as well as the inter-spike-interval histogram while rearranging the phase relationship between spikes and LFP. In summary, the 2D entropy of Φ and its correlation with navigation were immune to permutations of inter-spike-intervals, but both orderedness and behavioural correlation were significantly reduced by the phase-randomization of LFP (Extended Data 2.18 and Extended Data Table 4). The concordance of the two results argue that grid cell firing emerges from the underlying phase topography of LFP.

Next, we compared the gamma Φ and theta Φ with respect to entropy, low-spatial-frequency power and the 2D correlation of these maps with the avatar’s movements in space in terms of RVL. With the exception of otherwise high behavioural correlations (Kruskal-Wallis test *p*=0.4528, *r*_(Φ,*RV L*)_>0.5 *p*_(Φ,*RV L*)_<0.0001 for both theta and gamma Φ-s), gamma Φ displayed a significantly more ordered structure (smaller entropy) and larger low-frequency components than theta Φ (Kruskal-Wallis tests 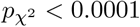 Extended Data Fig. 5d and Extended Data Table 3).

To tease apart the contribution of single neurons to the population phase, we compared the Φ across different electrodes as well as across individual neurons isolated from the same electrode. First, we isolated 16 single units from 4 different electrodes (4 neurons per electrode) and investigated the phase composition of *ϕ*-s and Φ-s by combining cells at electrode-level. Phase was expressed relative to gamma and theta LFP recorded form a separate electrode (Fig. 3a-f). The *ϕ* histograms displayed a highly polarized phase distribution (Rayleigh p<0.001; Fig. 3a). Despite the moderate gridness in FR maps, the gamma Φ expressed strong periodicity in alignment with the environment (Fig. 3a). When comparing the phase angles between electrodes, we observed a 60° rotation of phases between electrode 1 and its nearest neighbours, electrode 2 and 5. We also observed a 120° phase difference between electrode 1 and 3 (second neighbour). Notably, the observed phase tuning was specific to the gamma band LFP (Rayleigh p<0.001 Fig. 3a-f). The 80° difference between the two modes of the bimodal phase tuning was independent from electrode position. On one hand, the constant 60° rotation of phase tuning over equidistant electrodes suggests a constant 10 ms phase delay of gamma peaks between adjacent electrodes, consistent with the gamma autocorrelation function in Figure 1c and inter-electrode-delays of gamma (Extended Data Figs. 1 and 2). On the other hand, the rotation-invariant 80° bimodal phase tuning suggests that neurons maintain a constant phase relationship with the propagating gamma field. Moreover, the gamma phase of combined single unit activity displayed a significant dependence on the avatar’s heading RVL in all electrodes (χ^2^ test p<0.05), but no similar relationship was observed in theta phases (χ^2^ test *p*=0.1157) (Fig. 3 sixth column).

Finally, we asserted the contribution of individual neurons to the phase composition of a group of neurons recorded from the same electrode (Fig. 4b-e). The *ϕ* of single-units isolated from the group displayed a highly polarized phase tuning with five equiangular petals (Fig. 4b under *ϕ*). The other two neurons displayed only two components of the equiangular phase tuning, one with 144 degrees and another with 72 degrees between spokes (Fig. 4c,d *ϕ* and Φ) all in co-registration. The highly polarized *ϕ*, typical of gamma LFP, was in stark contrast with the lack of phase tuning with respect to theta (Fig. 4e). Despite the sparse spatial clustering owing to the low firing rate of cells (~1 *spike* * *s*^−1^), iso-phase nodes maintained their association with locations across cells (Fig. 4a-e Φ). Although theta *ϕ* was homogeneously distributed across the spectrum (Rayleigh *p*=0.62757), it displayed a clear dependence on the RVL of heading (χ^2^ test *p*_(*n*=196)_ = 7.3819*e*^−6^; Fig. 4e *ϕ*-heading). While single-units expressed evenly distributed phase tuning they did not correlate or cluster with the RVL of movement (Fig. 4a). In summary, different cell-assemblies sampled by different electrodes display a phase rotation consistent with a travelling gamma waves. At the same time, neurons near each other keep their discrete phase tuning in co-registration with each other and relative to the local gamma, while the same neurons display a dispersed distribution relative to theta. Although single neurons did not express gamma phase dependency on behaviour as they did in group (Fig. 3), the same single unit as featured on Figure 4a did express a theta phase dependency on the RVL of heading (Fig. 4e; χ^2^ test *p*=7.3819e-06, n=196).

Phase coherence^14–16^, including phase precession^17–20^, between action potentials and LFP has been studied in a great detail within and across cortical areas in rodents and primates, including humans^9,21^. However, because all these studies investigated the spike phase in a linear maze^20^ or during omnidirectional passes^22,23^, the reconstruction of spike phases was never complete.

By comparing the phases of the same spikes relative to gamma and theta band LFPs, we found that the correlations between virtual locations and spike phases were 3-10 times larger in gamma than in theta, and gamma phase was tuning showed more polarity (Figs. 3f,4e) and iso-phase nodes displayed shorter average inter-node distances (Extended Data Figs. 6k, 8) than theta, contrary to earlier efforts focusing on theta^13,22,23^. Gamma phase tuning and the power of low-frequency components in the spectral density of autocorrelograms computed from gamma phase maps were larger than those of theta phase maps. Moreover, the entropy of gamma phase maps was smaller than that of the theta phase maps (Extended Data Fig. 5d,e).

The second main observation was that the coefficient of variation of population-wide gamma phase modulation was smaller than firing rate grids’ (Fig. 1g,h,j). This suggests that spike phase is able to encode information under extremely low or high firing rate conditions, when gridness from firing rate is not evident. This observation was supported by the immunity of phase maps to ISI shuffling while phase randomization of LFP compromised the spike phase maps significantly (Extended Data Fig. 5b).

The third observation was that the inter-electrode phase shift was consistent with the model of propagating gamma waves^15,24^ that made the activity of cells recorded from two adjacent electrodes displaying a constant phase difference. We also observed, that gamma phases from single neurons as well as from multiple neurons recorded from the same electrode resulted in multi-modal phase distributions (Fig. 4b).

Lastly, we showed that neither gamma nor theta phase was independent of the spatial behaviour in our subjects. We demonstrated that the resultant vector of heading covaried with spike phases from the observed LFP more than with spike phases from phase-randomized LFP (Extended Data Fig. 5a), also consistent with the validation of phase patterns by surrogates with respect to entropy and low frequency spatial structures (Extended Data Fig. 5b,c). Phase coordination was found to rely on the LFP phase rather than the temporal statistics of spiking.

In summary, beyond phase tuning, we demonstrated a 2D modulation of spike-gamma phase coupling that is

(i) persistent, (ii) scales with the environment, and (iii) allocentric. The spatial scaling of phase modulation implicate that grid cells acquire their spatial specificity from the underlying field of gamma oscillations.

## METHODS

### 1. MATERIALS AND METHODS

#### 1.1 Subjects

Two male epilepsy patients (ages 35-41; average 38 years; (Table S1), previously consented, were implanted with microelectrode arrays in their EC in preparation for surgical resection of epileptic foci. From these two patients (subject 1 and 2) we could record well-isolated single unit activity with one channel local field potential (LFP) throughout a 7 and an 8-day period in the hospital’s epilepsy monitoring unit, while they performed a virtual navigation task that required spatial memory on a tablet computer (Extended Data Table 1).

#### 1.2 Tasks

The subjects’ were asked to play a computer game on a tablet they held on their lap. The game’s objective was to locate randomly dispersed space aliens in four different environments and return them to preassigned spaceships parking at memorized locations. The four virtual environments included a backyard (BY), a courtyard of the Louvre (LV), a model reconstruction of the main hall of the Luxor Temple in Egypt (LX), and a large open space (OS) with a boundless horizon and minimal external cues^4^. These environments differed in several features, including scenery, size, aspect ratio, and presence of obstacles or boundaries (Extended Data Table 2). The virtual reality environments were designed using Unity 3D (version 3.5.6.) and were compiled for Android 4.0. The game rendered the 3D environment from the player’s point of view. The player was constrained to the flat ground surface of each map and their movement speed was a constant 5 m/s, unless the “GO” button was released or an obstacle inhibited the movement.

#### 1.3 Game control

The task was performed on a tablet PC (ASUS Transformer 201 running Android 4.0 at 1280 × 800-pixel resolution). Subjects manoeuvred by pressing a “GO” button with their left thumb and controlled direction by pressing either a “LEFT” or “RIGHT” button with their right thumb. Before experimental data was collected, subjects were allowed to practice playing the game until they were accustomed to the game controls. The subjects’ virtual trajectory and heading (relative to the N-S axis in each game environment) were recorded.

#### 1.4 Implementation fo the task

Our subjects performed virtual spatial navigation tasks implemented as video games using a tablet computer, for a 5 or 10 minutes duration per game in 4 different environments (see Methods 1.2), total 40 minutes per day. The four environments differed in size, geometry, architecture, indoor, outdoor space and the richness of spatial landmarks^4^. The subjects viewed these scenes from a first person point of view (further referred to as the ‘avatar’s view’). The avatar’s movement was controlled by touch-screen buttons allowing movement controls of advancing along a straight or a curved trajectory by pressing GO and/or LEFT or RIGHT buttons at the same time, respectively. The touch screen control enabled also stopping (when not pressing GO) and turning while stopping. The avatar moved by emulating real walking with a constant step size and constant speed. The objective of the game was to locate randomly placed space alien, pick them up one by one, and deliver them to one of the two spaceships parking at constant locations of the space. The game program kept track of the avatar’s movements with a 16 ms sampling rate synchronized with the display frame rate. The pick up and delivery of a space alien was displayed on the screen giving a continuous feedback to the subject on his/her performance. We motivated the subjects to exceed his/her last day performance. Our subjects were able to complete as many as 50 space alien deliveries per day, average 2 space alien per minute.

#### 1.5 Synchronizing spatial navigation with neuronal data logging

The subjects’ navigation data, recorded on the tablet, was associated with the neuronal data by sending a 25 ms duration frequency modulated waveform from the tablet’s audio output port to the analog-auxiliary input port of the data acquisition system each time the “START” button for the game was released and periodically afterword. The precision of data synchronization between the tablet and the neuronal data logging was < 2 ms (SD 1 ms). This resulted in a spatial localization error of less than 2.5 cm virtual distance (< .048 % of average map width).

#### 1.6 Surgical procedures and electrode implantation and deplantation

We recorded wide-band signals from no deeper than layer II-III (given the < .8 mm tissue penetration and the average 5 mm cortical thickness of human EC, though lacking histological verification) of the medial entorhinal cortex. AD-TECH macro/micro subdural electrodes (Catalogue code: CMMS-22PX-F478), custom made per our specifications, were surgically implanted in the right hemisphere of two patients. The macro/micro electrode assembly consisted of 6 macroelectrodes and 16 microelectrode wires arranged in a 4×4 grid between the macroelectrodes. The microelectrodes were made of 35 um platinum iridium wires arranged in a 4×4 wire grid with 1 mm spacing between nearest electrodes. Electrodes were cut to .8 mm length from the electrode base and with nominal impedance < 3 *M*Ω. The craniotomy and electrode implantation were performed under general anaesthesia. After craniotomy, the electrodes were inserted subdurally to the surface of the entorhinal cortex by the neurosurgeon with stereotactic control. The dura was hermetically closed in a watertight fashion and the bone flap was reattached. The patient remained in the hospital ICU under continuous epilepsy monitoring for 5 to 14 days following the surgery. After sufficient evidence for seizure origin had been collected, electrode explantation and surgical resection of the seizure foci were performed under general anaesthesia.

#### 1.7 Recording neuronal data

Simultaneous single unit activity was obtained from five out of 16 microelectrodes at 24 kHz sampling frequency using an FHC Guideline 4000 system, an FDA approved amplifier for neuronal data acquisition in the human brain. The 5 electrodes varied across the days and were selected before the recording session based on the largest amplitude and most promising single unit isolation. The 5 or 10 minutes traces were band-pass filtered (300 to 6,000 Hz) by using a non-causal elliptic filter off-line. Because we selected 5 out of 16 electrodes with the highest unit activity each day before data logging we are unable to claim identity of single units across different days. Simultaneous LFP was sampled from all the 5 electrodes filtered digitally by a non-causal *filtfilt* filter between.1 and 300 Hz.

#### 1.8 Spike detection

We applied WaveClus off-line spike detection and spike sorting^25^. Spike detection was followed by isolation of single-unit activity using an unsupervised spike-sorting method. For spike detection we applied a threshold fitted to the median standard deviation of the data (1):

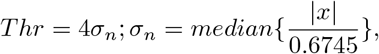

where x is the band-pass filtered signal and *σ*_*n*_ is an estimate of the standard deviation of the background noise. In cases when the amplitude threshold did not provide a clear separation between single and multiunit activity, the multiunit activity generated a large “noise cluster” in the wavelet coefficient space at near zero amplitude. This isolated noise cluster enabled us to separate single unit clusters from noise with high confidence. We only included single unit activity in our dataset if it was separated from the noise cluster by d>20, where d is the Mahalanobis distance^4^.

#### 1.9 Spike sorting

Spikes different from noise were sorted using the WaveClus method that uses superparamagnetic clustering as a non-parametric classifying engine^25^. WaveClus is the second most popular semi-supervised method worldwide, used by more than 110 publications and the most efficient among the bench-marked spike sorting methods^26^. The wavelet transform is defined as the convolution between the signal x(t) and a Haar wavelet functions *ψ*_*a,b*_(*t*),

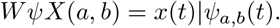

where *ψ*_*a,b*_(*t*) are dilated (contracted), and shifted versions of a unique wavelet function *ψ*(*t*),

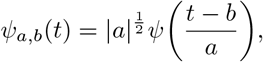

where a and b are the scale and translation parameters, respectively. Finally, we obtained 12 wavelet coefficients and reduced those to 4 dimensions with the highest multimodality and deviation from normal distribution. These were the dimensions best discriminating the spikes in wavelet coefficient space. Each spike was associated with a combination of these k most informative wavelet coefficients, hence represented by a point in the k-dimensional space. The data using superparamagnetic clustering resulted in clusters associated with spikes of similar waveform, where k=12 for all present datasets. The spike times of classified wave-forms were tested against 4 ms refractoriness before associated with putative neurons. We only included neurons where the Mahalanobis distance between the centroid of the noise cluster and the elements of single unit cluster or between the centroid of a given single unit clusters and the elements of other cluster was d > 20. We refer to these single unit clusters as activity of putative ‘neurons’ (Fig. 1a). Spike times were rounded to the nearest 1 ms interval and expressed in 1 ms precision.

## 2. COMPUTATIONAL METHODS

### 2.1 Characterizing the avatar’s movement in space

The subject’s task in the game was to navigate the avatar to pick up randomly displaced space aliens and reach memorized targets with them. During the course of navigation, some areas were visited more often than others resulting in an inhomogeneous distribution of sampling of the environment. The 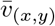 expressed the distribution of direction crossing at (x,y) area of the environment. The map of directions 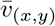 and the resultant vector length 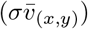 quantified the avatar’s movement in 4 dimensions as the X-Y location, the mean heading direction 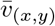 (angle) and the resultant vector length of heading direction 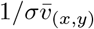 (the inverse of the variance of directions), given that the avatar was moving with a constant speed.

### 2.2 Computing heading direction and resultant vectors

The X-Y coordinates of the avatar’s movements and heading directions in the environments were up-sampled to 1 kHz by cubic-spline interpolation to match with the temporal resolution of neuronal data. Data synchronization was achieved through audio trigger pulses generated by the tablet and recorded through an analog-auxiliary input of the data acquisition computer (for method details see^4^). From the positions and facing directions (in angular degrees) we constructed a probability density of visits over each unit area of an environments during each game. For the construction of these maps we divided each environment uniformly by a square grid that was proportional to the size of environments (.7×.7 m for the small ‘BY’, 2×2 m for the large ‘LX’, ‘LV’, and 3×3 m for the largest ‘OS’ environments). Next, we determined the total amount of time spent in each square area, the number of crossings in that area, the circular mean direction of movement 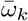 from a set of vectors representing the avatar passing through at a given *ij* unit area and the resultant vector of all passes. The length of the resultant vector served as an estimate of the consistency of heading directions over an area. Conversely, the inverse of the resultant vector length described the variance of directions of passing. The average direction 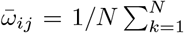 was defined as the mean direction of vectors at a given unit area. The mean direction informed us how stereotypical the view was from that location and the resultant vector length (RLV) informed us directly about the variance of the directions the avatar took by crossing the place. Given the average constant speed of the avatar, a small resultant vector *m*_*ij*_ indicated a large variance of directions, while large *m*_*ij*_ implied consistent directions. Direction and resultant vectors were all normalized by the number of vectors in the area. Four types of correlations with phase of spikes were computed: (i) Circular correlation between phase angles and motion direction angles^27^; (ii) Circular to linear correlation between the spike phases and RVL; and (iii) independence of spike phase and motion direction distributions vectors using χ^2^ tests, and (iv) independence of spike phase and RVL distributions using χ^2^ tests.

### 2.3 Circular to linear correlation between spike phase and movement resultant vectors

To correlate average neuronal activity (firing rates or spike phases) with movement parameters (heading direction or resultant vector) at any spatial location, we counted the spikes and trajectory segments over small areas each environment was divided into. All trajectory segments crossing a given area was aggregated and average direction and resultant vectors were computed. Likewise all spikes within that area were aggregated and mean circular phase of spikes were computed. Next, we computed the correlation between mean spike phases and the resultant vector length as circular to linear correlation^27^. If our data consist of *n* pairs of movements velocity (*m*_11_, *m*_12_, *m*_1*n*_) and spike phase angle (*a*_21_, *a*_22_, *a*_2*n*_), the circular correlation is defined^28^.

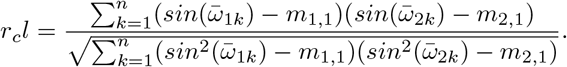

### 2.4 Correlation between movement direction and spike phase

Let assume our data consist of *n* pairs of movement angular velocity (*ω*_11_, *ω*_12_, *ω*_1*n*_) and spike phase angle (*ω*_21_, *ω*_22_, *ω*_2*n*_). The circular correlation is defined^28,29^.

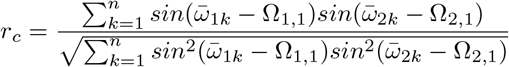

where Ω_1,1_ and Ω_2,1_ are the grand mean direction of the movements and spike phases, respectively. The estimated *p* value associated with the correlation is based on the assumption that *z*_*r*_ is distributed as standard normal

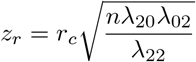

and

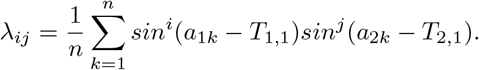

### 2.5 Binning the space for firing rate maps and phase maps

To elucidate the spatial aspect of spike phase (*ϕ*), the navigation arena was divided into uniformly sized (1×1 m or 2×2 m or 3×3 m) squares (unit areas) proportional to the total area of the environment. The size of the unit area did not influence the results within a range of .5×.5 m to 3×3 m^4^. For each of those areas we determined the times of visiting and the number of spikes generated inside the area. The computation of firing rate maps and phase maps is described under 2.12 and 2.13, respectively.

### 2.6 Computing 2D entropy

Instead of using a traditional grid-score metrics^30,31^ we used entropy for comparing phase maps of original spike trains with surrogate spike and LFP processes. The advantage of entropy over gridness is that (i) it is more general than gridness (ii) less sensitive to specific features such as rotational symmetry, (iii) it is agnostic to the grid distance and rotation symmetry unlike gridness, while it is sensitive to the periodic structure of the 2D image, (iv) easy to interpret and straightforward to compute. If *p*(*z*_*i*_) is the grey-level histogram of the phase map then the entropy of the image is^32^:

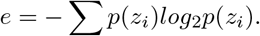

### 2.7 Analysis of grid parameters

Firing rate maps, spatial autocorrelograms (ACs), autoperiodograms were computed using standard methods^4,30,31,33,34^. We quantified grid scores by precisely following the method outlined by Barry et al, Sargolini et al, and Krupic et al^30,31,33^. Grid distance was determined based on the autoperiodogram and manually cross-validated with the autocorrelograms. To compute confidence intervals for statistical significance of gridness scores, we applied a standard Poisson bootstrap method and shuffled spike times 1,000-times. Validation of spatial periodicity against the by-chance was done using a Monte Carlo method by comparing the spectral modulation depth of each autocorrelogram against the distribution of gridness scores of 1,000 randomized autocorrelogram generated from mixtures of 2D Gaussian distributions. Not to confuse the Poisson bootstrap method for grid score validation with the inter-spike-interval shuffling applied for the validation of spike-LFP phases (see Methods 2.17 below).

### 2.8 Computing the firing rate grid period (grid distance)

Grid period is the wavelength of the spatially periodic single unit activity. It is equivalent with distance between adjacent nodes of the autocorrelogram. Since autocorrelograms are periodic by construction, this spatial wavelength is defined as the inverse of the predominant spatial frequency component, also could be measured by hand as the average grid distance^1^. Grid distances were measured following the method outlined in Nadasdy et a.^4^. Briefly, after the removal of the central peak from the AC (non-specific to the spatial pattern) we computed the 2D spectral density of the ACs by taking the complex conjugate of the inverse 2D Fourier transform^31^. We next averaged the 2D spectral distribution across the X and Y coordinates and determined the largest amplitude peak positions. The peak position corresponds to the predominant spatial frequency component of the grid. This method was chosen because it is more precise and less biased than measuring the distance between the nodes by hand. Dividing the dimensions of the AC by the spatial frequency provided the distance of the X-Y peak in spatial bins. We then computed the Euclidean distance of the peak (defined by its X and Y coordinates) from the origin, the centre of the autoperiodogram. This distance was multiplied by the scalar bin size (in m) to give the main grid period λ. Grid frequencies were computed for ACs generated by each neuron and compared between environments. Not to confuse the firing-rate grid distance with the “grid-phase node distances” (described in the next paragraph: 2.7 Computing grid phase node distance).

### 2.9 Computing grid iso-phase node distance

In the lack of a standard method for quantifying the spatial distribution of spike phases projected onto a 2D plane, we applied a manual method. First we plotted the colour-coded phase maps for every single unit that exceeded 1 Hz average firing rate, providing n>=300 spikes during a 5 minutes navigation session. The justification for the 1 Hz firing rate threshold was empirical as it provided the sufficient coverage for generating continuous phase gradients and discernible ISO-phase nodes, i.e. areas where the same phase repeats. We calibrated each phase map according to the size of the virtual environment and digitized the position of iso-phase nodes relative to the edges of the environment. Defining iso-phase nodes started with dividing the phase spectrum to four equal segments (1 – 90°,91 – 180°,181 – 270°,271 – 360°) corresponding roughly with blue, red, yellow and green colours of the HSV colour-map. Next we asked unbiased volunteers to mark the centroids of areas on the phase maps where one of the four colours is represented by at least 3 connected pixels and enter the coordinates into separate spreadsheets. To avoid a bias, the volunteers were blinded to the purpose of the study and the goal of measurements. Once the phase maps were digitized, we computed the Euclidean distances between each nodes (N) of the same colour and determined the inter node distances between them. The total number of distances within an iso-phase node graph is 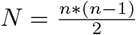. Next we constructed the distribution of these distances. For a periodic graph the distribution of inter-node distances formed several prominent peaks with sub-harmonics. The first peak in the distribution provided the average nearest neighbour inter-node distance (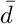. The mean of 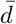 across the four phase range (colour) was used to express the iso-phase distance (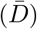.

### 2.10 Datasets and statistical methods

To compare grid scores and grid periods across environments, the general linear model ANOVA and its non-parametric version the Kruskal-Wallis test (Matlab, MathWorks^®^, Nattick, MA) were applied. The main factor was the environment (BY, LV, LX, OS), and the dependent variables were grid period and phase inter-node distance. We performed Rayleigh tests for testing non-uniformity of circular data and Watson’s goodness of fit test for testing conformity with the von Mises distribution (Matlab^®^ Circular Statistics Toolbox^27^).

### 2.11 Processing local field potentials (LFP)

Wide band signals were recorded from all 5 electrodes, but we computed the phase of single unit activity relative to LFP recorded from only dedicated electrode, referred to as El 5. For this electrode, we down sampled the original wide-band recording from 24 kHz to 1 kHz and digitally filtered the LFP between 0.1 to 300 Hz for broad spectrum overview by *filtfilt* a non-causal, zero-phase digital filter implemented in Matlab (https://www.mathworks.com/help/signal/ref/filtfilt.html). LFP was then further filtered at specific frequency bands by also using *filtfilt* for theta (1-12 Hz) and gamma (25-35 Hz). These frequency intervals were determined based on the prominent frequencies of the FFT (Fig. 1b). We selected epochs for the phase analysis when the spectral density function at theta or gamma band deviated by more than 2 * STD from the 1/f regression. Hence the number of epochs included in the theta and gamma phase analysis differed.

### 2.12 Computing spike to LFP phase

To get a precise phase estimate of the spikes, we sampled the spike waveform at 24 kHz sampling rate. We determined the spike time based on the largest first derivative of the positive going component (associated with the *Na*^2+^ influx) of action potentials. This time point was then rounded to the nearest 1 ms scale and associated with the LFP also sampled at 1 kHz. Next, to obtain the instantaneous phase, the Hilbert-transforms of both theta and gamma frequency filtered signals were computed using the hilbert function of Matlab (https://www.mathworks.com/help/signal/ref/hilbert.html). The Hilbert-phase at spike times (both defined at 1 ms precision) served as instantaneous phase estimate of spike relative to the theta or gamma oscillation.

### 2.13 Spatial firing rate maps

The spatial tuning of single unit activity was characterized by firing rate maps (FR maps) and their spatial autocorrelation functions. For the firing rate map, the game area was binned into 1×1, 2×2 or 3×3 m unit areas, depending on the size of the virtual environment (Extended Data Table 2) For each of those unit areas we determined the duration of time spent and the number of spikes generated during crossings. By normalizing the number of spikes by the time spent we obtained the firing rate (FR) in spikes/s (Hz). Binning had no significant effect on the grid parameters^4^).

### 2.14 Computing phase maps

Besides the local firing rates, the circular average phase of spikes (*ϕ*) relative to the theta and gamma band-pass filtered LFP was computed. Because the frequency of visits varied area-to-area, including no visit at all, the reliability of (*ϕ*) estimate also varied with the number of visits. Therefore, we computed the mean and variance (*σ*^*ϕ*^) of the phase estimate for each area. By spatially integrating the local phase estimates we constructed a global map 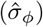, where the mean phase estimate 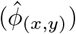 at any given area was associated with a colour of the HSV colour cylinder (red= −π, green=0, and red=π), while 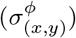 was represented by the value (*σ*_*max*_ = 2π = black and *σ*_*min*_ = 0 = maximum hue) (Fig. 1j,l). We refer to the variance-weighted spatial distribution of a phase plot as ‘phase map’ denoted by (Φ).

### 2.15 Computing autocorrelation (AC)

To compute the autocorrelation, the firing rate map was first smoothed with a Gaussian filter (5×5 bin neighbourhood, (*σ* = .8) and non-visited bins, originally assigned with NaN, were replaced by firing rate = 0 *spike* * *s*^−1^. Autocorrelograms were computed as follows. Given that the original firing rate map is *f* and the number of overlapping bins between the original and shifted firing rate maps at a given *τ*_*x*_, *τ*_*y*_ offset is *n*, the equation for the two-dimensional discrete autocorrelation is as follows:

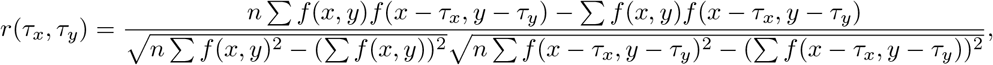

where *r*(*τ*_*x*_, *τ*_*y*_) is the autocorrelation. Correlations were estimated for all values of *n*. The central peak of the autocorrelogram was removed before computing the gridness^34,35^.

### 2.16 Computing grid scores

We quantified canonical “gridness” based on the autocorrelograms (ACs) by computing a 60° gridness score (g) step-by-step following the exact procedure outlined by Barry et al^30,31,33^ as described in our earlier study^4^. We first normalized the firing-rate maps by the sizes of environments that allowed for equal spatial resolutions for the ACs of different environments, but we kept the aspect ratio differences. Next, we computed the 2-dimensional ACs by applying 2D cross-correlation to the firing-rate maps^4^. After centring and clipping the AC to a 100 × 100 matrix, we located the largest peak after the removal of central peak, which defined a concentric ring containing the circular or ellipsoid arrangement of the first set of autocorrelation peaks at radius *R*. The outer radius of the ring was, based on the BK method, chosen to be 2.5*R*^4,30,31,33^. For the computation of gridness scores we followed the method by Sargolini et al^33^. Accordingly, we filtered the AC with the above-defined ring. Then we rotated the extracted ring from 1 to 180°, and computed the Pearson’s correlation coefficients *r*̄_(1°… 180°)_ between the original and rotated matrix with an 8-point moving average applied to it. We determined gridness g as the difference between the minimum of *r*_60°_ or *r*_120°_ and the maximum of *r*_30°_, *r*_90°_ or *r*_150°_. This function of gridness assumed a 60° modulation of AC as it expresses the modulation depth relative to 60° rotation symmetry. Because r-modulation was limited between 0 and 1, g was bounded between 0 and 1, inclusive.

### 2.17 Generating surrogate spike-trains

For testing the deterministic spike-LFP phase relationship against by-chance phase coincidences we generated surrogate spike-trains. To preserve the inter-spike-interval (ISI) statistics of the original spike train, yet decouple spike times from the phase of LFP, we re-sampled the inter-spike intervals from the ISI histogram and distributed them randomly during the interval of the LFP. We refer to this surrogate as ‘ISI-shuffling’. The first ISI of the original spike train was considered relative to time 0. ISI-shuffling provided a NULL to test the topographical consistency of spike-LFP phase relationship. We reasoned if the observed spike-LFP phase coupling is topography preserving then randomizing the phase relationship, while retaining the statistics of both ISIs and LFP, should lead to a dispersion of topography. Phase topography preservation was tested by cross-validation (see Methods 2.20 below). Even if spike trains were highly periodic (with a narrow ISI histogram) and potentially increase the by-chance phase-coupling between spikes and LFP, the ISI-shuffling would retain that by-chance coupling. In addition, to test whether or not periodic spike processes are sufficient to model the observed spike to LFP phase relationship, we created spike-trains with uniform inter-spike-intervals by determining the average inter-spike interval (1/f) and distributed the same number of spikes evenly within the recorded interval. Because trajectories were controlled by the non-deterministic placements of navigation targets, we can rule out any systematic or periodic coupling between the avatar’s positions and spike processes. Hence, the periodic spike processes alone cannot account for topography-preserving spike-LFP phase relationship.

### 2.18 Generating surrogate LFP

Testing the deterministic contribution of LFP to the observed spike to LFP phase relationship, we generated LFP surrogates by phase randomization of the LFP. We applied a phase decomposition of the original LFP, which preserved the original power spectral density, except the phases of oscillatory components were randomly shifted relative to the original. If spike processes were coordinated with the phase of any LFP component, that relationship was dissolved by phase randomization. As a result, the observed topographic structure of spike to LFP phase relationship should have been compromised in the surrogate. In order to remove the low frequency coherence between spikes and LFP we reversed the phase randomized LFP in time. As a result, the phase randomized and time-reversed LFP should destroy any systematic phase relationship between spikes and LFP.

### 2.19 Comparing phase maps of original to surrogate spike and LFP

We compared the original (Φ) with those of constructed from surrogate spike-trains and surrogate LFP with respect to (i) correlation of phase *ϕ*_(*x,y*_) and variance of heading direction 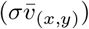 (the inverse of resultant vector length) (ii) grid entropy, and (iii) the power of the low-frequency components of the autocorrelation of (Φ). Statistics summarized in Extended Data Table 3. While random LFP phases significantly decreased the covariance of local spike-phase with the resultant vector length (*zval*^*subj*−1^=10.5241, *p*_(*n*=910)_=0.0, *zval*^*subj*−2^=7.8262 *p*_(*n*=1004)_=0.0), ISI-permutation did not (*zval*^*subj*−1^=−0.6207, *p*_(*n*=910)_=0.5348, *zval*^*subj*−2^ =0.2321 *p*_(*n* =1004)_=0.8165)(Extended Data Fig. 5a). Similarly, random LFP phase increased the entropy of (Φ), while ISI-permutation did not (*zval*^*subj*−1^ =0.3204, *p*_(*n*=910)_=0.7487, and *zval*^*subj*−2^=0.9118, *p*_(*n*=1004)_=0.3619) (Extended Data Fig. 5b). Moreover, the original (Φ) patterns displayed a significantly larger low-frequency power of the 2D-Fourier transform of phase maps than the ISI-shuffled and phase-randomized LFP (ISI-shuffling: *zval*^*subj*−1^=2.6879, *p*_(*n*=697)_=0.0072, *zval*^*subj*−2^=11.8777 *p*_(*n*=1004)_=0.0 and phase-randomized LFP: *zval*^*subj*−2^=8.6097 *p*_(*n*=1004)_=0.0), except subject 2 for LFP phase randomization (*zval*^*subj*−1^=1.6668, *p*_(*n*=697)_=0.0956) (Extended Data Fig. 5c).

### 2.20 Testing the consistency of phase topography by cross-validation

To test for temporal stability we split a 300 s spike and LFP data into two non-overlapping 150 s duration epochs and computed an element-to-element correlation between the phase maps (Extended Data Fig. 4a-d). We excluded the non-visited areas from the correlation that would otherwise generate spurious correlation. For the 2D cross-correlation, we applied Pearson’s correlation between the two vectorized maps following 2D cross-correlation formula:

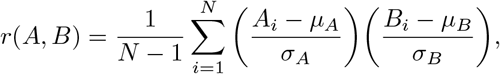

where *r*(*A,B*) is the cross-correlation, A and B were matrices representing the two phase maps, *μ*_*A*_ and *σ*_*A*_ are the mean and standard deviation of A, respectively, and *μ*_*B*_ and *σ*_*B*_ are the mean and standard deviation of B. For circular correlation the same formula was used as above except (*ω*_11_,*ω*_12_,*ω*_1*n*_) and (*ω*_21_,*ω*_22_,*ω*_2*n*_) are the gamma phases of corresponding elements in the two matrices *μ*_*A*_ and *μ*_*B*_ representing the means, and *σ*_*A*_ and *σ*_*B*_ were the variance of A and B matrices, respectively^28^.

For testing the robustness of the spike-gamma phase association, we eliminated the odd numbered spikes first, the even numbered spikes second, constructed phase-maps A and B (Extended Data Fig. 4a,e-g). Next, we computed the element-wise 2D correlation coefficients between the two matrices, alike we did to the split-half dataset above.

### 2.21 Generating autocorrelograms of surrogate spike to LFP phases

The topographic maps of spike to LFP phase relationship is affected by the inhomogeneous coverage of space by the time-limited navigation, no matter we were using the original spike trains and LFPs or we were using the surrogate counterparts. All reflected the pattern of spatial coverage, hence increased the correlation between original and surrogate patterns (path-correlation). To remove the path-correlation confound when evaluating difference between surrogate and original spike and LFP processes, we constructed autocorrelograms between the spike-LFP phase maps. The non-visited areas of maps ware rendered by random phase values, hence resulted low average correlations, while visited areas reflected true correlations between original spike/LFP and surrogate spike/LFP. We constructed autocorrelograms of true spike to LFP phase maps, surrogate spike to true LFP phase maps, and true spike to surrogate LFP phase maps.

## References

1. Hafting, T., Fyhn, M., Molden, S., Moser, M.-B. & Moser, E. I. Microstructure of a spatial map in the entorhinal cortex. Nature 436, 801–806 (2005).

2. Ekstrom, A. D. et al. Human hippocampal theta activity during virtual navigation. Hippocampus 15, 881–9 (2005).

3. Jacobs, J. et al. Direct recordings of grid-like neuronal activity in human spatial navigation. Nature neuroscience (2013). doi:10.1038/nn.3466

4. Nadasdy, Z. et al. Context-dependent spatially periodic activity in the human entorhinal cortex. Proceedings of the National Academy of Sciences of the United States of America 114, (2017).

5. Lu, C. B., Jefferys, J. G. R., Toescu, E. C. & Vreugdenhil, M. In vitro hippocampal gamma oscillation power as an index of in vivo CA3 gamma oscillation strength and spatial reference memory. Neurobiology of Learning and Memory 95, 221–230 (2011).

6. Buzsáki, G. Theta oscillations in the hippocampus. Neuron 33, 325–40 (2002).

7. Bragin, A. et al. Gamma (40-100 Hz) oscillation in the hippocampus of the behaving rat. J Neurosci 15, 47–60 (1995).

8. Colgin, L. L. et al. Frequency of gamma oscillations routes flow of information in the hippocampus. Nature 462, 353–357 (2009).

9. Rutishauser, U., Ross, I. B., Mamelak, A. N. & Schuman, E. M. Human memory strength is predicted by theta-frequency phase-locking of single neurons. Nature 464, 903–907 (2010).

10. Heusser, A. C., Poeppel, D., Ezzyat, Y. & Davachi, L. Episodic sequence memory is supported by a theta-gamma phase code. Nature Neuroscience (2016). doi:10.1038/nn.4374

11. Yartsev, M. M., Witter, M. P. & Ulanovsky, N. Grid cells without theta oscillations in the entorhinal cortex of bats. Nature 479, 103–7 (2011).

12. Killian, N. J., Jutras, M. J. & Buffalo, E. A. A map of visual space in the primate entorhinal cortex. Nature 491, 761–764 (2012).

13. Jacobs, J. Hippocampal theta oscillations are slower in humans than in rodents: implications for models of spatial navigation and memory. Philosophical transactions of the Royal Society of London. Series B, Biological sciences 369, 20130304 (2014).

14. Vinck, M. et al. Gamma-phase shifting in awake monkey visual cortex. The Journal of neuroscience: the official journal of the Society for Neuroscience 30, 1250–7 (2010).

15. Besserve, M., Lowe, S. C., Logothetis, N. K., Schölkopf, B. & Panzeri, S. Shifts of Gamma Phase across Primary Visual Cortical Sites Reflect Dynamic Stimulus-Modulated Information Transfer. PLOS Biology 13, e1002257 (2015).

16. Siegel, M., Warden, M. R. & Miller, E. K. Phase-dependent neuronal coding of objects in short-term memory. Proc Natl Acad Sci USA 106, 21341–21346 (2009).

17. O’Keefe, J. & Recce, M. L. Phase relationship between hippocampal place units and the EEG theta rhythm. Hippocampus 3, 317–330 (1993).

18. Jones, M. W. & Wilson, M. A. Phase precession of medial prefrontal cortical activity relative to the hippocampal theta rhythm. Hippocampus 15, 867–873 (2005).

19. Skaggs, W. E., McNaughton, B. L., Wilson, M. A. & Barnes, C. A. Theta phase precession in hippocampal neuronal populations and the compression of temporal sequences. Hippocampus 6, 149–172 (1996).

20. Hafting, T., Fyhn, M., Bonnevie, T., Moser, M. B. & Moser, E. I. Hippocampus-independent phase precession in entorhinal grid cells. Nature 453, 1248–1252 (2008).

21. Jacobs, J., Kahana, M. J., Ekstrom, A. D. & Fried, I. Brain oscillations control timing of single-neuron activity in humans. J Neurosci 27, 3839–3844 (2007).

22. Climer, J. R., Newman, E. L. & Hasselmo, M. E. Phase Coding By Grid Cells In Unconstrained Environments: Two-Dimensional Phase Precession. European Journal of Neuroscience 38, 2526–2541 (2013).

23. Reifenstein, E., Stemmler, M., Herz, A. V. M., Kempter, R. & Schreiber, S. Movement dependence and layer specificity of entorhinal phase precession in two-dimensional environments. PLoS ONE (2014). doi:10.1371/journal.pone.0100638

24. Bahramisharif, A. et al. Propagating Neocortical Gamma Bursts Are Coordinated by Traveling Alpha Waves. Journal of Neuroscience 33, 18849–18854 (2013).

25. Quiroga, R. Q., Nadasdy, Z. & Ben-Shaul, Y. Unsupervised spike detection and sorting with wavelets and superparamagnetic clustering. Neural Comput 16, 1661–1687 (2004).

26. Wild, J., Prekopcsak, Z., Sieger, T., Novak, D. & Jech, R. Performance comparison of extracellular spike sorting algorithms for single-channel recordings. Journal of Neuroscience Methods 203, 369–376 (2012).

27. Berens, P. CircStat: A MATLAB Toolbox for Circular Statistics. Journal of Statistical Software 31, (2009).

28. Jammalamadaka, S. R. & SenGupta., A. Topics in circular statistics [electronic resource]. 322 (World Scientific, 2001).

29. Fisher, N. I. & Lee, A. J. A correlation coefficient for circular data. Biometrika 70, 327–332 (1983).

30. Barry, C. & Bush, D. From A to Z: a potential role for grid cells in spatial navigation. Neural systems & circuits 2, 6 (2012).

31. Krupic, J., Burgess, N. & O’Keefe, J. Neural Representations of Location Composed of Spatially Periodic Bands. Science 337, 853–857 (2012).

32. Gonzalez, R. C., Woods, R. E. & Eddins, S. L. Digital Image Processing Using Matlab - Gonzalez Woods & Eddins. 624, 609 (2004).

33. Sargolini, F. et al. Conjunctive representation of position, direction, and velocity in entorhinal cortex. Science 312, 758–762 (2006).

34. Migliore, M. et al. ModelDB: making models publicly accessible to support computational neuroscience. Neuroinformatics 1, 135–9 (2003).

35. Hines, M. L., Morse, T., Migliore, M., Carnevale, N. T. & Shepherd, G. M. ModelDB: A Database to Support Computational Neuroscience. Journal of computational neuroscience 17, 7–11 (2004).

